# BeeSAM2: detecting bees in cherry flowers using timelapse images and foundational models

**DOI:** 10.1101/2025.08.01.668108

**Authors:** Jane Devlin, Fraser MacFarlane, Alison Karley, Fabio Manfredini, Dominic Williams

**Affiliations:** University of Aberdeen; Janes Hutton Insitute; James Hutton Institute

## Abstract

Bees perform important pollination services in fruit crops such as cherry. Growers will often bring in bees to supplement natural pollinators. Monitoring the performance of these pollinators is important to understand the impact of augmenting pollinators on fruit yield particularly in relation to June drop which is a major cause of yield instability in the cherry industry. Timelapse imaging plus automated image analysis methods is a valuable tool in studying the role of bee pollination in fruit set. Timelapse cameras allow for continuous monitoring of the flowers, but manual analysis of the generated footage is very time consuming.

We have developed a novel method of detecting bees in time lapse images, called BeeSAM2. This exploits both the zero shot detector *Grounding Dino* and the foundational model *Segment Anything* 2. Promising results are achieved with the method able to detect the bumblebee *Bombus terrestris* in images with a recall of 0.959 and precision of 0.991. These results are accurate enough to deploy our method to quantify bee activity in cherry plantations, advancing the ability of researchers to monitor bee interactions with flowers with a significant time saving over manual analysis of timelapse footage.

## Introduction

Pollinators are very important for crop production worldwide. In commercially grown sweet cherry plantations, self-sterile cherry varieties are often grown, meaning that pollen from another compatible variety carried by bees or other insects is needed for successful pollination. Cherry trees suffer from a phenomenon known as “June Drop” where immature fruit is dropped by a tree. The drivers of this process are not fully understood but it is thought that it may be related to pollination (Gatti 2024) (Mir, Mir et al. 2025). In particular the quality of pollination services such as visitation rate, duration, stigma contact and flower handling may all affect both initial fruit set and subsequent fruit drop. This has motivated the study of pollination in cherry trees.

There is a variety of methods that can be used to monitor pollination by bees: these include trapping pollinators, walking transects, eDNA, and the use of cameras (Howard, Nisal Ratnayake et al. 2021, Johnson, Katz et al. 2023, Van Klink, Sheard et al. 2024). The use of cameras to monitor pollinator behaviour is an increasing area of interest (Steen 2017, Ratnayake, Dyer et al. 2021); the advantages of using cameras include allowing precise monitoring of flowers without requiring to be in the field for extensive periods of time and lower pollinator disturbance compared to a human observer. Researchers have used a combination of timelapse photography and computer vision methods to analyse these type of data. Due to the small size of bees and wind movement of flowers the use of motion sensor cameras presents many challenges so continuous recording and post capture analysis of footage is used instead.

Several studies have used timelapse photography combined with automated detection methods to monitor insects. (Ngo, Rustia et al. 2021) used a camera system to monitor honey bees entering and leaving a bee hive. They used a simple USB camera connected to a raspberry pi which continuously recorded bees at the entrance of the hive. Bees were classified according to whether they were carrying pollen or not using a model based on YOLOv3 (Redmon and Farhadi 2018), characterized by tiny architecture and achieving high accuracy (0.91 precision, 0.99 recall) for detection of bees with pollen loads. (□tefan, Workman et al. 2024) used smart phones mounted on a tripod to monitor pollinator interactions with flowers. They recorded timelapse footage at 1second intervals for one hour. No automation was used for image analysis and over 1,720 hours were spent labelling the 460,000 gathered images indicating the high time cost associated with analysing data if computer vision techniques are not used. (Bjerge, Frigaard et al. 2023) used timelapse images of flowers to monitor insect visitation primarily by honey bees. They set up recording units containing USB web cameras connected to a raspberry pi looking at flowers recording images at 30s intervals. Images were captured from 4:30h to 22:30h. They used a combination of motion enhancement and YOLOv5 (Jocher, Stoken et al. 2020) object detection method to automatically detect insects in the images. The model was trained on 3,783 images of which 2,499 contained insects. The same group used a similar system without the addition of motion enhancement instead aiming to classify different insect pollinators detected again using YOLOv5 (Bjerge, Alison et al. 2023).

One of the challenges in using object detection methods is the requirement for specific training and a frequent lack of generalisability with the models produced. This has prompted interest in producing “foundational” models which are able to generalise to new problems without the need for finetuning. Segment anything model (SAM) (Kirillov, Mintun et al. 2023) aims to segment any objects in an image going from either point or bounding boxes to indicate the object. By prompting systematically across an image, it can detect all objects in an image. It has been used extensively in various imaging problems including medical image analysis (Zhu, Hamdi et al. 2024, Sengupta, Chakrabarty et al. 2025), crop image analysis for the detection of leaves (Williams, Macfarlane et al. 2024), detecting size of cocoons in images of bee colonies (Getz, Best et al. 2024) and to help segment images of moths (Jain, Cunha et al. 2024). Segment anything model 2 (SAM2) (Ravi, Gabeur et al. 2024) is an advancement on SAM which aims to track objects through video frames. An initial prompt is required which is used to select an object of interest and then this is followed through all the frames in a video. It achieves state of the art results on several standard video segmentation benchmarks.

Alongside foundational models, there has also been research into zero shot object detection methods, i.e. object detection that doesn’t require training. Grounding dino (Liu, Zeng et al. 2024) is an example of this type of method. A text input is given and matching objects in an image are detected. This method has been used extensively to solve a variety of problems. (Mullins, Esau et al. 2024) tested different text prompts using both grounding dino and YOLO-World for a series of tasks on blueberry plants. They found grounding dino outperformed YOLO-World for most tasks and required fewer descriptive prompts finding blueberries with prompt “smooth blueberries” rather than “a small blue sphere”. Grounding dino has been used to help crop images of parasitoid wasps prior to taxonomic identification (Shirali, Hübner et al. 2024). Grounding Dino and SAM have been combined by (Ren, Liu et al. 2024): here grounding dino was used as a zero-shot object detector and SAM was used to generate a segmentation of this object.

So far limited work has been done to incorporate zero shot detectors or foundational image models to the task of insect pollinator detection. In this paper we present a novel method that combine grounding dino and SAM2 to detect bumblebee pollination events in timelapse images of cherry flowers: this method has the potential to be applied to other bees or pollinating insects in different scenarios.

## Materials and methods

### Plants

The experiment was carried out in two polytunnels with two rows of cherry trees in each. The trees were arranged in blocks of the same variety. Image data was gathered from the variety Kordia which was present in both tunnels. Kordia is a self-incompatible variety so needs to be cross-pollinated with a different variety to develop fruit. In tunnel 1, three varieties, Kordia, Sweetheart and Regina, were grown in 5 plant plots, with a total of 45 trees in each row. Sweetheart and Regina are both compatible with Kordia for pollination. In the second tunnel, four varieties, Kordia, Penny, Lapins and Sweetheart were grown in 4 plant plots. Penny, Lapins and Sweetheart are all compatible pollinators for Kordia. At the time of the experiment the trees were 6 years old. Three commercial colonies of the bumblebee *Bombus terrestris audax* were added to the tunnels to provide pollinators for the cherry flowers, mimicking standard commercial practice by cherry growers in Scotland. Insect netting was used to create three sections only accessible to commercial colonies (one colony per section) and three sections that were open to wild pollinators to enter: each section contained 15 trees, 5 each of Kordia, Sweetheart and Regina. 12 Afidus timelapse ATL-200S cameras (two per section) were mounted on wooden posts and pointed towards a single flower cluster on the target tree. The trees were randomly selected but each flower cluster was selected to fit criteria of being approximately 1.5m from the ground, at approximately the same stage of development and be in a recordable position. They were set to take timelapse photos at a rate of 1 per second between 11am and 5pm, corresponding to peak foraging times for commercial bumblebees: this was done to reduce the recording time with the aim of extending the battery life of cameras.

### Manual analysis of timelapse images

The cameras stored the timelapse images as video files. These were then watched by an observer and any visits of bees to the target flower cluster was recorded with information on time of arrival and departure, duration, bee species, and flower identity. The purpose of this annotation was to record pollination events, so only bees that landed on the target flower clusters were recorded; bees which either failed to land on flowers or landed on flowers outside of target cluster were not recorded. This was a very time consuming process – approximately 500 hours which motivated the development of an automated method to complete it more quickly in future applications.

### Automated Bee detection methods

A number of methods were developed aiming to automatically detect bee visits from the timelapse footage with minimal human intervention. For all methods, the individual frames were extracted from the timelapse footage. Grounding dino was used, setting a prompt as “bee” for the target object with a confidence threshold of 0.3 used for any object found. We refer to this method in the results and discussion sections as GD_bee.

Further testing of additional prompts was carried out to determine the best prompt to use to detect bees in the images. Use of closely related prompts such as “wasp”, “insect”, “pollinator” produced similar results but slightly more missing observations. One of the challenges faced while detecting bees during a pollination event was occlusion caused by the bees landing on the flowers and being partially hidden within the flowers. To overcome this problem the prompt of “pollinating bees” was used to provide the model with additional context to improve results. Initial testing was also carried out for prompts of “bees in flowers” but any inclusion of flower resulted in detection of flowers without bees, so this was not continued. In the results and discussion sections this method is referred to as GD_pollinating_bee.

### Segment Anything Model 2

Using segment anything 2 (SAM2) requires definition of an object to track on the first frame of the video. In our case most of the frames did not contain bees. Therefore, a synthetic first frame was added to the beginning of the videos. This was created by taking the first frame of the video and then adding a bee taken from one of the subsequent images known to contain a bee. This bee was placed in the image 4 times (i.e. in the same image on 4 separate occasions) to enable tracking of multiple bees. An example of what this looks like is shown in figure1. The bee was flipped for two of the instances. SAM2 was then prompted with 4 different objects one for each bee. Once the initial object was selected, SAM2 attempted to track the object through all the other frames in the video, and it maintained the ability to detect objects even after they had not appeared in the video for 100s of frames. While the purpose of the method was to track unique instances of an object, we aimed to detect the presence/absence of bees rather than counting individual bees. Our testing indicated that SAM2 could not easily distinguish individual bees and so was suitable for use as a generic bee detection method. In the results and discussion this method is called BeeSAM2.

**Figure 1.**
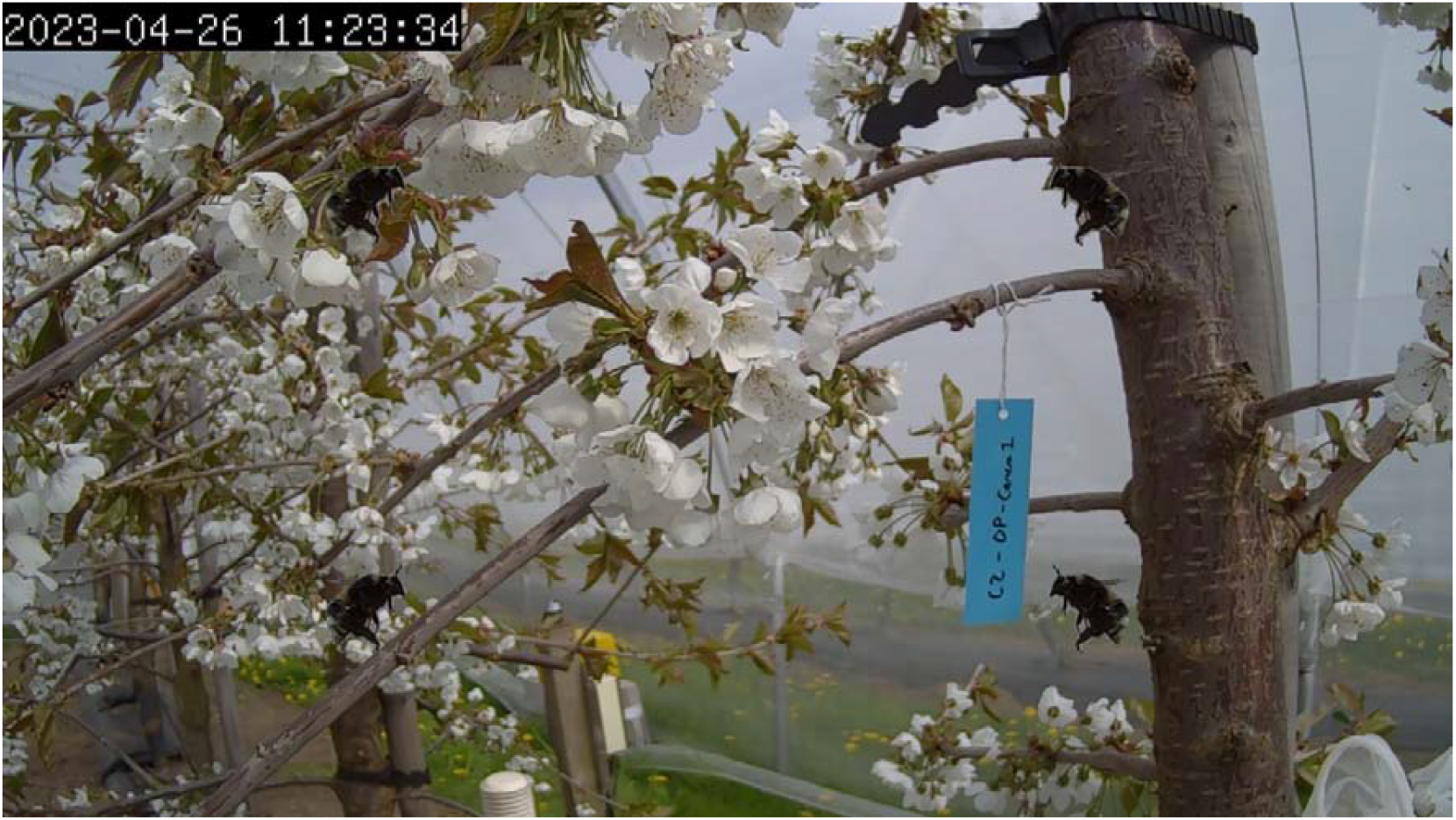
Example synthetic image with four bees added to it. The image was obtained by imposing the same picture of a bee in 4 different locations and 2 orientations on a background image representing a cluster of cherry flowers.

In order to improve the results of this method a combination of grounding dino and SAM2 was used. The first step was to run grounding dino through all frames in the video. The prompt of “bee” was used with a confidence threshold of 0.6. This was done to minimise the number of false positives. Any bees found by grounding dino were used to mark objects in the image for SAM2 to track. Frames in which no bees were detected were not labelled to avoid the risk of feeding false negatives into SAM2. The synthetic image was used for the first frame of the sequence. For all subsequent frames, labels from the bees detected by grounding dino were added. After labelling the bees in the images, the labels were propagated through the rest of the video using SAM2. In the results and discussion this method is known as GD_BeeSAM2.

### Evaluation of results – our data

To quantify the success of the different methods we used the manually analysed timelapse footage data. This was footage where the time and duration of bees landing on target flowers had been recorded by an observer. The exact location of the bee in the image was not recorded by the manual observer so it was assumed that any bees detected in the same frame where the observer recorded a bee on a target flower was a successful bee detection. The method developed here aimed to detect all bees, not just those on the target flowers, and so we produced an additional category of false positives: correct detection of bees that were not on target flowers. In order to quantify this component, an additional annotation was carried out for all the frames with a false positive bee detection recording, to discern whether they were truly a false positive or were a correct detection of a bee. Frames where no bees were detected were not checked again, so it is not possible to quantify the proportion of bees away from target flowers that were detected in the image (but not recorded manually).

We used the following measures to compare the different methods: recall, precision and adjusted precision. Recall is defined by the formula:

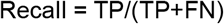

where TP is true positives – i.e., bees on target flowers accurately detected - and FN is false negatives - or number of bees on target flowers missed by model.

Precision is defined by the formula:

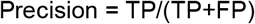

where TP is bees on target flowers accurately detected and FP is false positives – i.e., number of bees in frames where there are no bees on target flowers (so including accurate detection of bees off target flowers).

Finally, Adjusted Precision is defined by the formula:

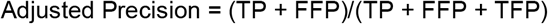

where TP is bees on target flowers accurately detected, FFP is bees detected on non-target flowers and TFP is frames where detected bees do not correspond to a bee i.e. another object was detected that was not a bee.

### Evaluation of results – data from (Bjerge, Frigaard et al. 2023)

To allow for a comparison to be made with previous methods we also evaluated our method using the dataset from (Bjerge, Frigaard et al. 2023), which consists of timelapse images with labelled bee objects. We again evaluated performance on a frame level assuming that any bee detected in the frame was the same as the labelled bee in the same frame. Our method includes the creation of a synthetic image for the first frame with four bees added to an image as described above. Two different methods were used: the first involved using the same bee as in our study and adding this 4 times to the first frame of new footage; the second consisted of finding a new bee from the Bjerge study using grounding dino and adding it to the first frame of each video (again 4 times).

## Results

The methods that we tested showed different measures of recall and precision (Table 1). Precision was low for all methods, with a max of 0.29 for GD_bee. This is due to the way we calculated precision and indicates the detection of many bees not located on the target flowers. We will instead focus on the adjusted precision scores as a more accurate reflection of the precision of the methods.

**Table 1:** Recall, precision and adjusted precision of the four versions of methods presented here. GD_BeeSAM2 has highest recall and adjusted precision.

|  | **Recall** | **Precision** | **Adjusted Precision** |
| --- | --- | --- | --- |
| GD_bee | **0.580376** | **0.288681** | **0.971942** |
| GD_pollinating_bee | **0.795876** | **0.062887** | **0.308583** |
| BeeSAM2 | **0.937213** | **0.130351** | **0.906699** |
| GD_BeeSAM2 | **0.958967** | **0.12933** | **0.990552** |

**Table 2:** Recall and precision of Grounded Bee Sam2 on data from (Bjerge, Frigaard et al. 2023). Note here there is no adjusted precision as they annotated all bees present in their video file.

|  | Recall | Precision |
| --- | --- | --- |
| GD_BeeSAM2 | **0.804** | **0.812** |
| GD_BeeSAM2_alternativebee | **0.831** | **0.832** |

GD_bee had the lowest recall of 0.58 but had a high adjusted precision of 0.97. This showed that grounding dino ‘out of the box’ can be used to detect bees with reasonable accuracy. However, with a recall of only 0.58 it could not be confidently used to determine if a flower had been visited by a bee.

Changing the prompt to ‘pollinating bee’ resulted in an increase in the recall score to 0.80: however this resulted in a significant reduction of the adjusted precision score to 0.31, indicating that most of the objects found using this method were not bees, limiting its usability as a method.

BeeSAM2 had better results with a recall of 0.94 and an adjusted precision of 0.91. This showed good potential for this method. However, once it started to track an incorrect object it would generally continue to track it unless there was further intervention. A large part of the reason for lower precision when compared to GD_bee was the tracking of a leaf cluster instead of bees in one of the timelapse videos.

GD_BeeSAM2 was the best performing method in terms of both recall (0.96) and adjusted precision (0.99). These scores imply that this method could reliably be used to monitor bee flower visitation with good confidence that not many bees would be missed. It is worth noting that in some of the frames where a bee was known to be on the flower, the bee was very heavily occluded and the presence of the bee was only known to the manual observer by watching the sequence of frames and seeing the flower position shift by the weight of the foraging bee.

Figure 2 shows the location of all detected bees in the image. It demonstrates the reason for the use adjusted precision score, many of the false positives are clustered round off target flowers indicating that they were likely to be correct identification of bees.

**Figure 2.**
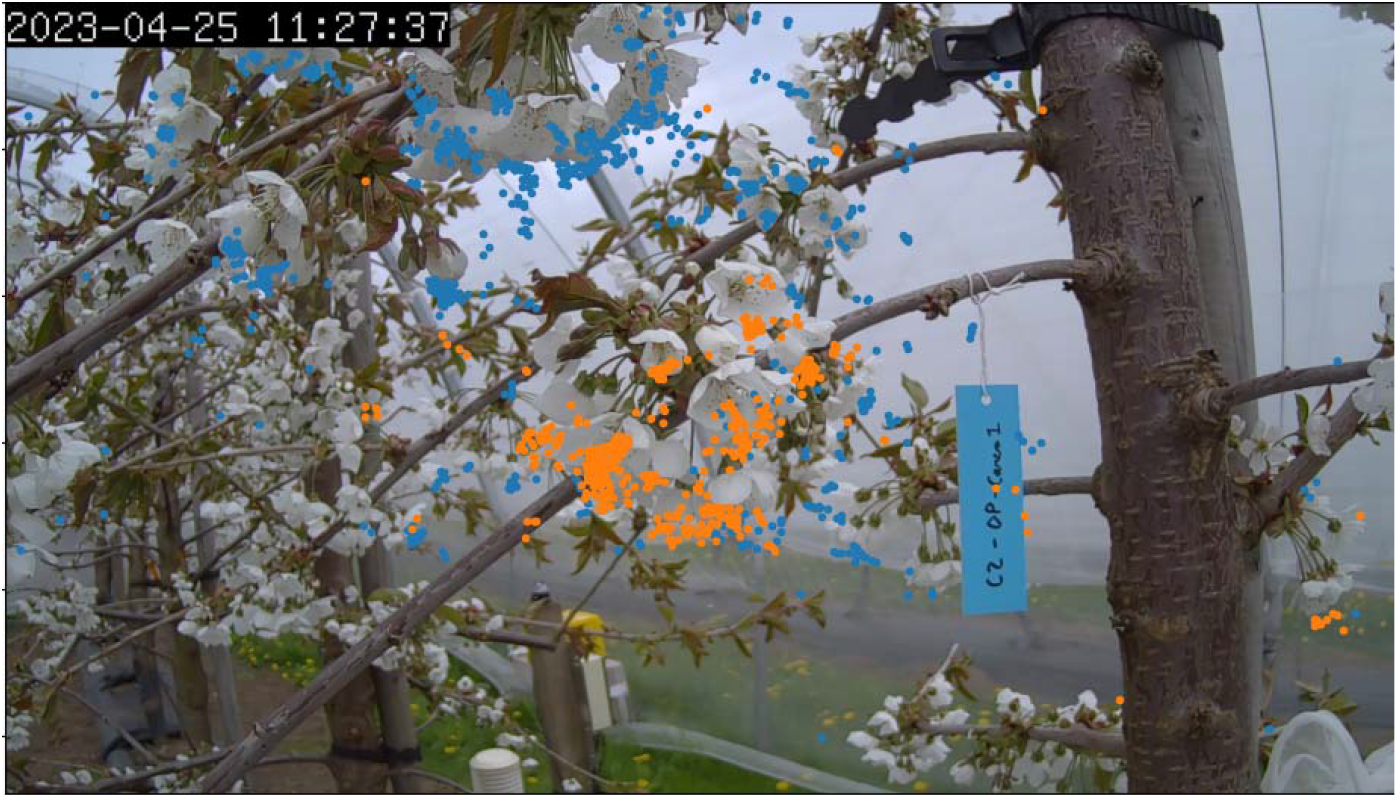
Locations of bee detections marked on an example image. Orange dots represent true positives and blue dots represent false positives. Target flowers cluster is at the centre of the image, bee visits on flowers in the background were not recorded. We can see here that many of the blue points are clustered around off-target flowers indicating that they may correspond to actual bee detections.

### Failure case examples

To understand the limitations of the methods proposed, it is useful to show some of the failure cases. One of the weaknesses of SAM2 is that it has no knowledge of the nature of the object it is tracking beyond the initial identification. Images a and b in figure 3 are an example of where a small bee on a flower was initially correctly identified, but in the following frames the tracking was transferred from the bee to the flower it was on. The method was initially designed to be interactive so this could be corrected with negative points, but of course this would not be suitable for fully automated detection of bees. Fine tuning of the model may be able to overcome this issue but would reduce the generalisability of the method described here.

**Figure 3.**
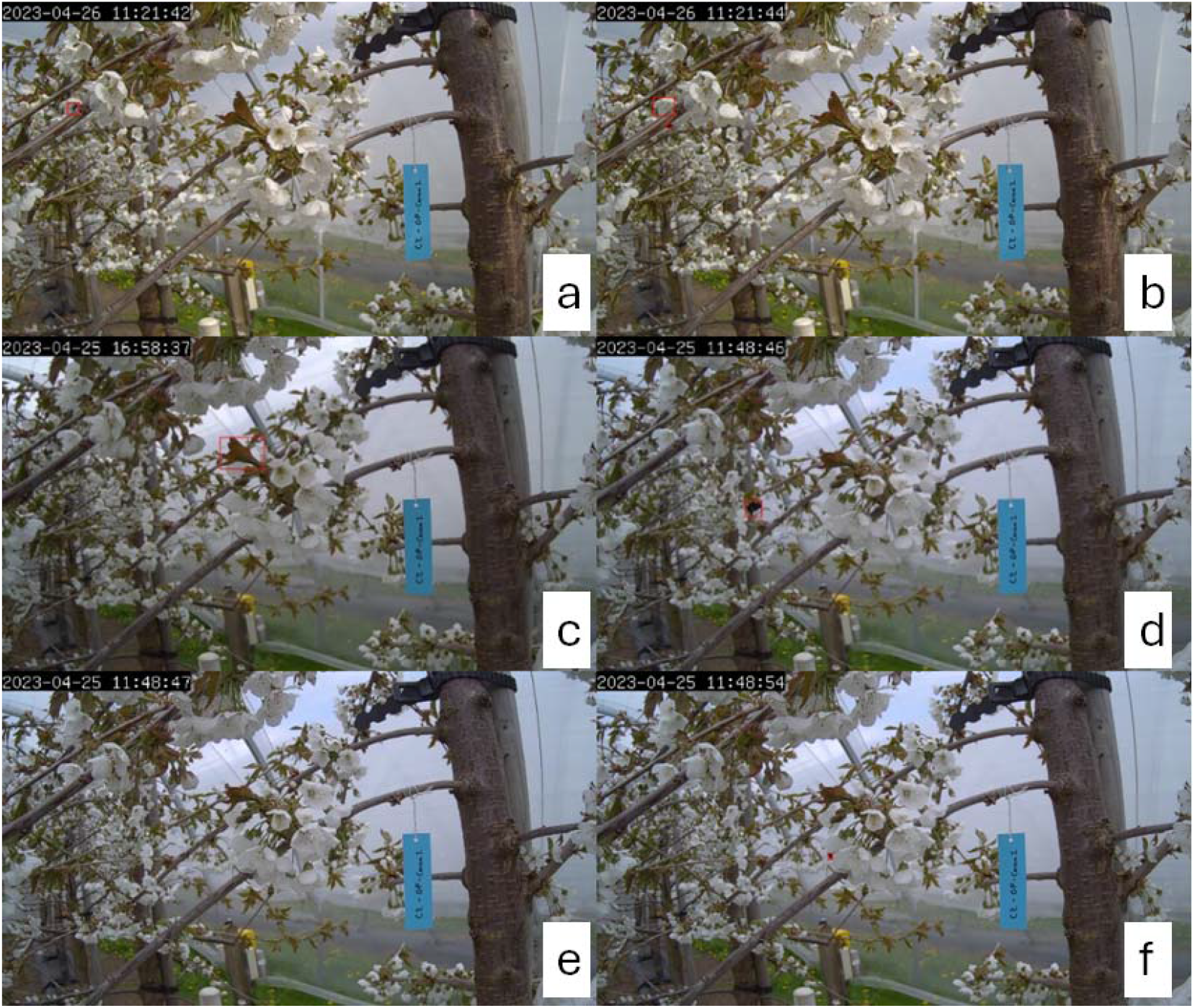
Example images showing failure cases. Images a and b show a sequence where a true positive detection (a bumblebee) becomes a false positive (a flower’s petal). Image c shows an example of a false positive caused by the false detection of leaf buds as a bee. The sequence d, e and f shows an example of a false negative due to a bee (initially rightly detected, panel d) becoming hidden within a flower in panel e and then re-emerging in panel f when exiting the flower.

Another failure case is shown in panel c in figure 3 (produced by Grounding dino). A few additional objects in the image were occasionally identified as bees - for example a cluster of leaves like shown in the picture. Once the prompt from grounding dino was given to SAM2 it would track that object indefinitely. In this case a single incorrect prompt from grounding dino was replicated for the rest of the frames of that video by SAM2. Increasing the confidence threshold used by Grounding Dino can help reduce the likelihood of this happening.

Looking at panels d, e and f in figure 3 we can see an example of a false negative. First, we can see a bee detected flying towards the flower (panel d); in the next frame (panel e) no bee is detected, as the bee has entered one of the flowers; finally, several frames later (panel f) the bee is detected again leaving the flower. If we manually observe the sequence, we can see that the bee is in the flower, so this sequence of frames was marked during manual classification as having a bee present; however, none of our models were able to detect this. Looking at several false negatives this appeared to be a consistent source of errors. This shows that while the model can detect some heavily occluded bees, it has some limitations compared to manual observers who can infer bee locations from a sequence in a way that our model cannot. This may cause problems for users trying to count how many bees are foraging on a specific flower, as it could result in considering a bee landing and then leaving the same flower as two separate visits.

### Assessment of our method with data from (Bjerge, Frigaard et al. 2023)

The application of the GD_BeeSAM2 method to data from (Bjerge, Frigaard et al. 2023) produced worse results than those observed for our dataset overall, but still achieved recall and precision greater than 0.8. A slight improvement in performance, from 0.812 to 0.832 for precision and from 0.804 to 0.831 for recall, was seen when using bees from the Bjerge et al dataset to generate the initial synthetic image. This shows that creation of a new synthetic image from the relevant data improves the performance of our method. These results are slightly worse but comparable to performance seen in the original paper where the authors achieved 0.888 recall and 0.897 precision on the same data set. The fact that our method closely matches the performance of a model trained specifically on the dataset highlights the potential of our model to generalise well and to be used on new unseen datasets.

## Discussion

In this study we have developed a method that is able to detect bees on flowers in timelapse video footage. The method detects bees with good accuracy, over 0.95 recall and 0.99 precision on new data that we obtained specifically for this study. Previous studies have reported much lower precision and recall using motion assisted YOLO based methods: for example, (Bjerge, Frigaard et al. 2023) achieved a recall of 0.89 and precision of 0.90. Their images included flowers from a range of different plant species and were recorded over a longer time period, so there may have been more variability in the data which could partly explain the relatively lower performance. The method described here, applied to the data from (Bjerge, Frigaard et al. 2023), performed worse than the Bjerge model itself, with a recall of 0.83 and precision of 0.83: however, it must be noted that these results were obtained without any training of the model on that dataset. In (Bjerge, Frigaard et al. 2023), the authors trained their model specifically on the data they used, requiring time consuming data labelling to be able to use the method. The method described here has no training or fine tuning required beyond the creation of a synthetic first image, making it generalisable to new settings easily.

When using our method on data from (Bjerge, Frigaard et al. 2023) we found that using bees from that study to create the synthetic image improved results. This indicates that further investigation into using 4 different bees for the initial synthetic image may improve results. Particularly if a wider variation of bee types were being investigated in a study. Given the very high recall and precision scores achieved on our data we did not feel the need to attempt this.

Other studies have focussed on monitoring the entrance and exit of honeybees to/from hives. In such studies, additional information on whether the bees were entering or exiting the colony and the presence of pollen baskets on bees was also required. Our model, as it stands, is not able to provide this information. Our output is a segmentation of the entire bee, with no information on the direction of movement. A second model, able to detect different parts of a bee, would be required to get additional information. However, considering that our model tracks bees through images, with sufficient video footage it may be able to track bees through frames of a video and therefore infer movement direction that way instead.

One potential addition to our model would be to add a classifier that can categorize bees based on whether they are in flight or have landed on a flower. This would help determine which flowers are undergoing pollination in the image. Nevertheless it is still possible to get a rough estimation of the two scenarios with our method, as in the output of our model bees on flowers will be detected multiple times, as they stay in the same place for longer, while bees in flight will appear in only a few frames. This can be seen in figure 2 which shows the location of detected bees. This approach could also be used to estimate the duration of the presence of a bee on a flower, a factor that seems to be positively correlated – until a certain point – with successful pollination.

## Conclusion

A method called BeeSAM2 has been developed to monitor bees on cherry flowers using timelapse imaging and automated image analysis. A high level of accuracy was achieved, showing that the method is suitable for use by scientists wishing to study bee behaviour in a cherry plantation. The model was not specifically trained on the data from our images, making it easily generalisable to new unseen data.

Unlike other previous studies, this method does not attempt to distinguish particular species of pollinators from each other. This is partly due to the trade off of using a generalisable model that can be applied to any species of insect, and also because in the experimental setup implemented to obtain the initial dataset, commercial colonies of *Bombus terrestris audax* were intentionally placed in the tunnels with the trees, meaning that the overwhelming majority of pollinators in the footage were *Bombus terrestris audax* workers.

## Author Contribution statement

*DW, AK, FM and JD designed the experimental setup; JD collected the data; DW and FM designed the methodology; DW and JD analysed the data; DW led the writing of the manuscript. All authors contributed critically to the drafts and gave final approval for publication*.

## Acknowledgments

The research was supported by The Anthony and Margaret Johnston Centre for Doctoral Training in Plant Sciences, Charles Sutherland Sponsorship travel grant, Internal Funding of The University of Aberdeen Grants Academy to Pump-Prime Research and Research Networks; Project Title: ‘Implementing the use of remote camera tools for the study of plant-pollinator interactions’, Scottish Society of Crop Research, and the Scottish Government’s Rural and Environment Science and Analytical Services Division (RESAS) through the strategic research program (2022-2027).

## Data Availability

Data is available here BeeSAM2: Image data

Code is available here Dom3442/BeeSAM2

